# The hydrolase LpqI primes mycobacterial peptidoglycan recycling

**DOI:** 10.1101/312876

**Authors:** Patrick J. Moynihan, Ian T. Cadby, Natacha Veerapen, Monika Jankute, Marialuisa Crosatti, Galina V. Mukamolova, Andrew L. Lovering, Gurdyal S. Besra

**Affiliations:** Institute of Microbiology and Infection, School of Biological Sciences, University of Birmingham, Birmingham, UK, B15 2TT; Department of Infection, Immunity and Inflammation, University of Leicester, Leicester, LE1 9HN, UK

## Abstract

Growth and division by most bacteria requires remodeling and cleavage of their cell wall. A byproduct of this process is the generation of free peptidoglycan (PG) fragments known as muropeptides. These muropeptides are recycled in many model organisms, where the bacteria can harness their unique nature as a signal for cell wall damage. These molecules also serve as important signals for hosts where binding to specific receptors reports on the presence of intracellular bacteria. Despite this critical role for muropeptides, it has long been thought that pathogenic mycobacteria such as *Mycobacterium tuberculosis* do not recycle their PG. Herein we show for the first time that *M. tuberculosis* and *Mycobacterium bovis* BCG are able to recycle components of their PG. We demonstrate that the core-mycobacterial gene *lpqI*, encodes an authentic NagZ β-*N*-acetylglucosaminidase and that it is essential for PG-derived amino sugar recycling *via* an unusual pathway. By characterizing an *M. bovis* BCG strain lacking *lpqI* we are also able to show that stem-peptide recycling proceeds independent of amino sugar recovery and loss of *lpqI* leads to antimicrobial resistance *in vitro*. Together these data provide a critical first step in understanding how mycobacteria recycle their peptidoglycan.

---

The cell wall of *M. tuberculosis* is built upon a foundation of peptidoglycan (PG). The remainder of this structure is formed by the modification of muramic acid residues with an arabinogalactan polymer that is in turn esterified by mycolic acids^1^. This waxy coating contributes to drug resistance in *M. tuberculosis*, but is also the target of several mycobacteria-specific antibiotics^1^. The challenge of multi- and extensively-drug resistant *M. tuberculosis* has not adequately been met by drug discovery efforts, however recent reports suggest that β-lactams are effective at treating these drug-resistant infections^2-4^. Despite their therapeutic promise, we know relatively little about the turn-over of PG in mycobacteria, which is the eventual target of β-lactam antibiotics.

For most bacteria maintenance of a PG sacculus is an essential aspect of life. PG is a heteropolymer comprised of glycan chains with a repeating disaccharide motif of *N-*acetylglucosamine β1 → 4 *N*-acetylmuramic acid (Glc*N*Ac-Mur*N*Ac) which are then cross-linked to one another *via* short peptides attached to the C-3 D-lactyl moiety of Mur*N*Ac (Figure 1a). The integrity of this macromolecule must be maintained under most growth conditions and its rupture leads to lysis and cell death^5^. As a result of this essentiality, it is vital that cells are able to withstand their own internal turgor pressure and still be able to cleave the cell wall to allow for division, growth and the insertion of macromolecular structures such as secretion systems^5^. Throughout this process, the activity of lytic enzymes or through the attack of host agents like lysozyme, the sacculus is cleaved with the resulting generation of small PG fragments^6^.

**Figure 1.**
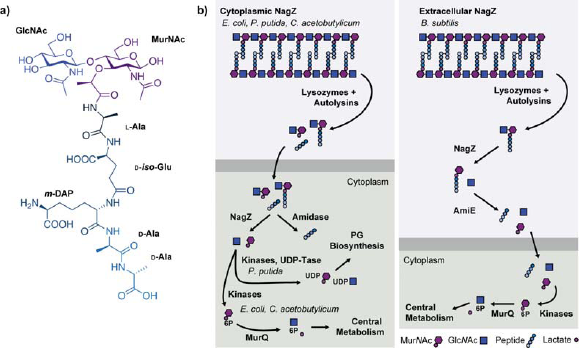
Overview of PG-recycling. **a)** The basic building block of PG is Glc*N*AcMur*N*Ac-pentapeptide. Enzymes produced by the bacterium or the host are able to cleave every major linkage in PG. **b)** The PG recycling machinery is variable with respect to the localisation of NagZ and the subsequent conversion to Glc*N*Ac-1P or UDP-Glc*N*Ac/Mur*N*Ac. All known Mur*N*Ac recovery systems that sustain bacterial growth (as opposed to strictly recycling e.g. *P. putida*) terminate at MurQ in the cytoplasm.

In Gram-positive bacteria muropeptides are typically released from the cell wall through the action of lysozyme-like hydrolytic enzymes, whereas in Gram-negative bacteria, lytic transglycosylases generate 1,6-anhydroMur*N*Ac products^7,8^. These metabolites have been shown to be important in many aspects of host-pathogen interactions. For example, tracheal cytotoxin produced by *Bordetella pertussis* is the product of lytic transglycosylases^9^. Release of a similar molecule has also been shown to be involved in tissue damage during *Neisseria gonorrhoeae* infection and in the closure of the light-organ of the bobtail squid^10,11^. In many organisms, soluble PG acts as a potent immune stimulator once sensed by NOD receptors and other pattern recognition receptors^12^.

Aside from host organisms, PG metabolites are also important signaling molecules for the bacteria themselves. Recycling of PG has been studied in great detail in a small number of organisms including *Escherichia coli, Pseudomonas aeruginosa* and *Bacillus subtilis* among others^13^. The recycling pathway typically involves the step-wise degradation of the polymer into its monomeric constituents, monosaccharides and amino acids (Figure 1b). Despite common biochemical steps, compartmentalization of these steps tends to be organism specific^7^. The resulting monosaccharides are eventually phosphorylated and Mur*N*Ac-6-phosphate is converted into glucosamine-6-phosphate through the activities of cytoplasmic MurQ and NagA enzymes (Figure 1b). At the same time, the stem peptides are degraded to smaller components and typically shunted back into PG biogenesis. Growth on Mur*N*Ac as a sole carbon-source has never been demonstrated for a bacterium that lacks MurQ. Furthermore, recycling of Mur*N*Ac in a bacterium that lacks MurQ has only been described in *Pseudomonas putida*, and many bacteria, including mycobacteria, are not thought to recycle their PG at all^14^.

In the present study we sought to determine if mycobacteria are capable of recycling their PG and if so, what impact this has on the bacterium. We for the first time reveal that these bacteria do indeed possess the biochemical capacity to recycle PG elements and determined the molecular basis of Mur*N*Ac recovery. Our data indicate that loss of a key recycling enzyme, LpqI, leads to increased antibiotic and lysozyme resistance.

## Materials and methods

### Bacterial strains and growth conditions

Unless stated otherwise all chemicals and reagents were purchased from Sigma Aldrich. *M. bovis* BCG (Pasteur) and related mutants were maintained on Middlebrook 7H10 agar or 7H9 broth supplemented with 10% OADC enrichment and 0.05% Tween 80. Where appropriate kanamycin or hygromycin was added at 25 or 50 μg•mL^−1^, respectively. *Mycobacterium smegmatis* mc^2^155 was maintained on Tryptic Soy Broth or Tryptic Soy Agar where appropriate. For growth on defined carbon sources, strains were cultivated in Sauton’s minimal medium containing per 1 L, 4 g asparagine, 2 g citric acid, 0.5 g K_2_HPO_4_, 0.5 g MgSO_4_ * 7 H_2_O and 0.05 g ferric ammonium citrate, 0.05% tyloxopol and 5 mM of each carbon source unless stated otherwise^15^. *Escherichia coli* strains were grown in lysogeny broth and supplemented with kanamycin at 50 μg•mL^−1^ or hygromycin at 150 μg•mL^−1^ where appropriate.

### Cloning and purification of Rv0237

Rv0237 was amplified from *M. tuberculosis* H37Rv genomic DNA using standard PCR conditions with the Rv0237SUMOF and Rv0237SUMOR primers and cloned into the TA site of the Champion pET-SUMO expression plasmid (Invitrogen) according to the manufacturers specifications (all primers are found in Table S2). For production of Rv0237 1 L of *E. coli* BL21 [pRv0237] grown in Terrific Broth to an OD_600_ of 0.6, chilled to 20 °C and induced with 1 mM IPTG and grown for a further 18 h before being collected by centrifugation. Cells were resuspended in 25 mM Tris-HCl, 300 mM NaCl, 10 mM imidazole pH 7.8 and lysed *via* three passages through a French pressure cell. The protein was purified using standard IMAC procedures with washes of lysis buffer, lysis buffer including 50 mM imidazole and finally eluted with 500 mM imidazole in lysis buffer. Eluted protein was dialysed exhaustively against 25 mM Bis-Tris, 100 mM NaCl pH 7.8 in the presence of recombinant Ulp1 protease which specifically cleaves the His_6_-SUMO tag. Digested protein was passed through a second IMAC column (1 mL HisTrap FF, GE Healthcare) and the flow-through fraction was found to contain pure, un-tagged Rv0237. Purified protein was dialysed into 25 mM Bis-Tris pH 6.5, 100 mM NaCl.

### Crystallography

Prior to crystallization, LpqI was concentrated to 20 mg•mL^−1^ in 25 mM Bis-Tris pH 7.5, 100 mM NaCl. LpqI crystals were grown by the sitting-drop vapour diffusion method by mixing an equal volume of protein solution with 1.1 M sodium malonate, 0.1M HEPES, 0.5% w/v Jeffamine ED-2001 (pH 7.0). Crystals were cryo-protected with a saturated solution of sodium malonate and flash -ooled in liquid nitrogen. X-ray data was collected at the Diamond Light Source, Oxford. Data were processed using XiaII and file manipulations were performed using the CCP4 suite of programs. The structure was phased by molecular replacement using the unpublished *M. smegmatis* LpqI structure (PDB: 4YYF) using the program PHASER. The structure was subsequently auto-built in PHENIX and the remaining parts were built in COOT with further refinement using PHENIX and PDB-REDO.

### Kinetic characterisation of Rv0237

Purified Rv0237 was evaluated for glycoside hydrolase activity using a variety of substrates. As an initial screening assay, Rv0237 was incubated at 1 μM with either 4-methylumbeliferyl or *p*-nitrophenyl derivatives of a variety of sugars as listed in Figure 4a in Bis-Tris pH 7.5, 100 mM NaCl at 37 °C. The release of *p*-nitrophenol was followed by change in absorbance at 420 nm while production of 4-menthylumbelliferone was monitored by fluorescence as above in a BMG Polarstar spectrophotometer. Kinetic characterisation of Rv0237 was conducted using varying concentrations of 4MU-Glc*N*Ac. The raw data were compared to standards of 4-methylumbelliferone. All data were analysed using GraphPad Prism 7.

To evaluate the ability of the enzyme to degrade fragments derived from PG, *M. smegmatis* PG was digested with mutanolysin and soluble fragments were prepared and quantified as above. Reactions including 1 μM Rv0237, 0.5 mM PG fragments in 25 mM ammonium acetate buffer pH 6.5 were incubated for 18h at 37 °C. In parallel reactions were carried out using *p*NP-Glc*N*Ac in order to monitor enzyme activity visually. The reactions were then evaluated by TLC (Silica 60 F_254_, Merck, Germany) using a mobile phase consisting of 1-butanol, methanol, ammonium hydroxide and water at a ratio of 5:6:4:1. TLCs were stained with α-naphthol and developed by charring.

### Mutant generation

To generate the Δ*lpqI* strain we used specialized transduction according to established protocols^16^. A recombinant *lpqI* knockout phage was designed to replace the chromosomal *lpqI* gene using homologous flanking regions to *lpqI* with a the *sacB* gene and a hygromycin resistance cassette in-between using the LL-Rv0237, LR-Rv0237, RR-Rv0237 and RLRv0237 primers. The resulting phage was transduced into *M. bovis* BCG and transductants were selected on 7H10-agar plates containing 75 μg•mL^−1^ hygromycin. The mutant was verified by PCR and phenotypically with 4MU-Glc*N*Ac where loss of *lpqI* was expected to abolish turn-over of this fluorescent substrate. The complemented strain was generated by incorporating the *lpqI* gene and 150 bp 5’ to the start codon containing the promoter sequence into the promoter-less integrative mycobacterial shuttle vector pMV306 using primers Rv0237CompF and Rv0237CompR to generate the resulting pMV306-*lpqI* plasmid^17^. This was electroporated into *M. bovis* BCG Δ*lpqI* and transformants were selected on 7H10 agar containing hygromycin and kanamycin. The complemented mutant was verified phenotypically with 4MU-Glc*N*Ac. A control strain was also generated using the empty pMV306 vector using the same protocols.

### Antimicrobial testing

Mid-exponential cultures of *M. bovis* BCG and derivative strains were diluted to OD_600_ = 0.1 in fresh 7H9 media. 100 μL of this culture was added to a 96-well plate with the addition of 1 μL of antibiotic/lysozyme to achieve the desired final concentration as indicated. These were incubated for 7 days at 37 °C at which point 30 μL of 0.02% w/v resazurin and 12.5 μL of 20% Tween 80 v/v was added to the culture. This was incubated over-night at 37 °C and the production of resorufin was determined by fluorescence (Ex. 530 nm, Em. 590 nm) using a BMG Polarstar plate reader.

### Rapid purification of mycobacterial cell wall

Rapid purification of cell wall from small cultures was carried out using a modified phenol extraction protocol^18^. Mycobacterial cells were grown to mid-exponential phase and collected by centrifugation. These were washed with cold phosphate-buffered saline (PBS) and resuspended in PBS and the cells were lysed in a Percellys Evolution Bead Beater at 5,000 rpm for 3 min. The lysate was then transferred to glass culture tubes to which 2 mL of 98% phenol was added and vortexed for 1 min. This was heated for 1h at 70 °C, allowed to cool and the insoluble material was collected by centrifugation at 3,220 x *g*. The aqueous phase was removed and 4 mL of methanol was added. This was vortexed and centrifuged again. Finally, the pellet was washed 3 times with methanol and once with water before being frozen or used for subsequent enzymatic treatment.

#### Large-scale purification of PG

Purification of PG from *M. smegmatis* was achieved following established protocols^19^. Six-liters of *M. smegmatis* were grown to mid-exponential phase (OD_600_ = ˜0.6) at which point they were harvested by centrifugation, resuspended in a minimal volume of PBS and lysed by sonication. The resulting lysate was brought to 4% SDS and boiled under reflux for 3 h. The insoluble material was collected by centrifugation and washed with water until the SDS was completely removed (at least 7 washes) to yield mycolyl-arabinogalactanpeptidoglycan (mAGP). The mAGP was incubated for 4 days in 0.5% KOH in methanol at 37 °C before being washed three times with methanol. The mycolic acids were extracted with 3 washes of diethyl ether. The phosphodiester linking the AG-PG complex was cleaved using 0.2 M H_2_SO_4_ and the PG was separated from the solubilized AG by centrifugation prior to neutralization with NaCO_3_ and washed with water 3 times. The insoluble PG pellet was sequentially digested with α-amylase (100 μg•mL^−1^), DNase (10 μg•mL^−1^) and RNase (5 μg•mL^−1^) for 8h before proteinase K (100 μg•mL^−1^) digestion overnight at 37 °C. The PG pellet was resuspended in a minimal volume of 1% SDS and boiled under reflux for 3 h before the SDS was removed by centrifugation and washing with water (at least 7 times). The resulting material was lyophilized and stored at −20 °C until it was needed.

Digestion of cell wall material with mutanolysin was carried out overnight at 37 °C in 20 mM ammonium acetate buffer (pH 6.0) with continuous mixing. Following digestion, solubilised muropeptides were isolated using graphitized carbon solid-phase-extraction cartridges as previously described^20^. Purified fractions were evaporated to dryness and the concentration of reducing sugars in the pool of soluble muropeptides was assessed using the 3-methyl-2-benzothiazolinone hydrazone (MBTH) assay^21^.

### Synthesis of 4MU-D-lactate

Instead of the 2- or 3- step protocols published for the synthesis of 4MU-D-lactate previously, we used a simplified one step method^22,23^. 1.5 g of (s)-(-)-bromopropionic acid was added to 1 g of 4-methylumbelliferone stirring in 40 mL anhydrous dimethylformamide and 0.75 g Cs_2_CO_3_. This was stirred at room temperature over-night and the product was extracted three times with water/ethyl-acetate and the organic phase was dried over sodium sufate. The organic phase was then filtered and evaporated to dryness. The product was subsequently purified using silica chromatography and was dried as a crystalline white solid.

### Turn-over of 4MU reporter compounds by*M. bovis* BCG

To test turn-over of 4MU-Glc*N*Ac or 4MU-D-lactate by whole cells, 100 µL of a mid-exponential culture (OD_600_ = 0.6) was added to a sterile 96 well plate in Sauton’s minimal media supplemented with 0.05% Tween and 1% glycerol in addition to 1 mM 4MU-D-lactate or 4MU-Glc*N*Ac. Similar controls lacking cells or the reporter compound were included as well. This was incubated at 37 °C and mixed at 300 r.p.m. Each day the fluorescence of the sample was read on a BMC PolarStar microplate reader (Ex. 355 nm; Em 460 nm) with a constant gain setting.

### Turn-over of *M. bovis* BCG PG *in vitro*

Cultures of *M. bovis* BCG wild-type, Δ*lpqI*, and Δ*lpqI*::*lpqI* were grown to an OD_600_ of 0.6 in the presence of 10 µCi ^3^H *meso*-diaminopimelic acid (DAP), at which point they were collected by centrifugation, washed 3 times with sterile media and diluted to 0.01 in fresh culture flasks. Periodically a sample of 0.5 mL was taken, and the cells were collected by centrifugation. The spent medium was mixed with 10 mL scinitilation fluid and counted using a liquid scintillation counter. The cell pellet was re-suspended in 10% SDS, boiled for 20 min, and centrifuged again. The cell-wall material was then resuspended in 1 mL scintillation fluid and the material was counted in a liquid scintillation counter. The counts of the cell wall and the media were added together to give total ^3^H DAP in each culture and the data is presented as a percentage of that total. During the course of the experiment the OD_600_ of the culture was monitored daily. All measurements are from three biological replicates.

## Results

### Peptidoglycan Recycling Genes in *Mycobacteria*

The genome of *M tuberculosis* encodes many lytic enzymes, including at least five Resuscitation Promoting Factors (Rpfs) and greater than 10 peptidases and amidases in addition to penicillin binding proteins with potential lytic activities^24^. The Rpfs are most likely lytic transglycosylases with the product of RpfB having been recently confirmed as a GlcNAc-1,6-anhydroMurNAc disaccharide^25^. While *M. tuberculosis* does appear to encode at least one lysozyme, Rv2525, its activity has not been demonstrated^26^. A recent comparative study of PG-active enzymes in mycobacteria indicated that while significant differences exist, enzymes that can likely degrade all of the major covalent linkages of PG are encoded in the genomes of all mycobacteria^24^. The products of most of these enzymes have not been experimentally demonstrated, however their conservation underscores the importance of PG-remodeling during growth and division of mycobacteria.

Most autolytic enzymes produce small PG-metabolites (muropeptides), indicating that mycobacteria should generate these molecules during the course of normal growth. Indeed, soluble PG fragment release has been observed for both *M. smegmatis* and *M. tuberculosis in vitro*^18,27^. Given the slow release of PG fragments by mycobacteria, we evaluated the presence of known PG-recycling systems in the genome of several corynebacterial species (Table S1)^14,28^. BLAST analysis of the *Corynebacterium glutamicum, M. tuberculosis, Mycobacterium leprae* and *Mycobacterium bovis* BCG genomes indicates that they lack genes related to any known muropeptide import proteins, PG-metabolite phosphorylation systems, and *murQ*. The only sugar-kinase orthologs identified in the genome have previously been characterized as glucose-kinases although they have not been directly tested for amino sugar-phosphotransferase activity^29^. This contrasts sharply with *M. smegmatis* for which an apparently complete “classical” muropeptide recovery system exists, making it a poor model for the PG metabolism of *M. tuberculosis* (Table S1). Taken together, the available data indicates that *M. tuberculosis* and almost all other mycobacteria lack most of the known PG-recycling genes from other bacteria, with only two conserved genes potentially associated with PG-recycling, (*nagA* – Rv3332, *nagZ* – Rv0237).

### Biochemical and structural characterisation of LpqI

In previously characterized PG-recycling systems free amino sugars are produced by NagZ, which belongs to the CAZy glycoside hydrolase family 3 (GH3)^30^. This family is a large group of enzymes which have hydrolytic and phosphorylytic activity and remove β-linked sugars from proteins and polysaccharides^31,32^. The β-*N*-acetylglucosaminidase sub-family including all known NagZ enzymes utilize a conserved Asp-His catalytic dyad which has been well characterized^33,34^. A BLAST search of the *M. tuberculosis* H37Rv genome revealed only one NagZ ortholog, which was previously named LpqI in light of its identification as a lipoprotein including an appropriately positioned lipobox at the N-terminus of the protein^35^. As a lipoprotein LpqI is expected to be found attached to the external face of the cytoplasmic membrane, which is consistent with proteomics results^35^. LpqI has also been identified as a likely mannosylated glycoprotein in a proteomics screen using ConA chromatography^36^. The *lpqI* gene is found in all mycobacteria with sequenced genomes including *M. leprae* which has a substantially reduced genome indicating that it is involved in a conserved process across all mycobacteria (Table S1, Figure S1).

Given the absence of other PG-recycling associated genes, we sought to identify the function of LpqI. While LpqI bears significant sequence similarity to known β-*N-*acetylglucosaminidases, recent studies have demonstrated that divergent activities for this sub-family of enzymes are possible^32^. These activities included the ability to release sugars other than Glc*N*Ac from reporter substrates and apparent phosphorolytic activity. To test this, we cloned, expressed and purified the LpqI protein from *M. tuberculosis* H37Rv using an N-terminal His_6_-SUMO tag which was subsequently cleaved from the protein. We first determined if the protein was in fact a β-*N*-acetylglucosaminidase by testing its activity on a variety of substrates including many sugars that would be found in the cell wall of mycobacteria. Using convenient reporter sugars we assessed the ability for the enzyme to release *p*-nitrophenolate or 4-methylumbeliferone from conjugated arabinose, galactose, galactosamine, arabinofuranose, glucose, mannose, mannosamine, glucosamine and *N-*acetylglucosamine (Figure 2a). While not exhaustive, this set of sugars covers most major modifications to the cell wall including the AG itself, *O*-mannose modifications of proteins, Gal*N* modification of arabinan, the rhamnose-linker sugar of AG and the Glc*N*Ac and Glc*N* found in PG. The only detectable activity for LpqI was with Glc*N*Ac-containing substrates. Critically, in mycobacteria this sugar is limited to the backbone of PG and a small amount in the linker unit (Mur*N*Ac-6-P-Rha-Glc*N*Ac-galactan) between PG and arabinogalactan. The Michaelis-Menten constants (*k*_cat_ = 2.8 × 10^−2^ ± 0.04 × 10 ^−2^ •s^−1^ and K_m_ = 106 ± 5 µM) of LpqI using 4MU-Glc*N*Ac as a substrate were found to be similar to other NagZ enzymes using this substrate (Figure 2b)^33^. In a similar assay we were also able to show that LpqI releases Glc*N*Ac from soluble PG fragments (Figure 2c). While hydrolytic activity has been reported for most NagZ-type enzymes, a recent report suggested that β-*N*-acetylglucosaminidases from the GH3 family are in fact phosphorylases^32^. Another GH3 β-*N*-acetylglucosaminidase was recently reported to lack this activity, suggesting that it may not be a general property of the family^37^. We tested the activity of the enzyme under the same conditions as reported previously for Nag3 from *Celulomonas fimi* and found that there was no detectable difference with our observed hydrolytic activity. The product of the reaction also co-migrated with Glc*N*Ac on TLCs and not Glc*N*Ac-1-P (Figure S2).

**Figure 2.**
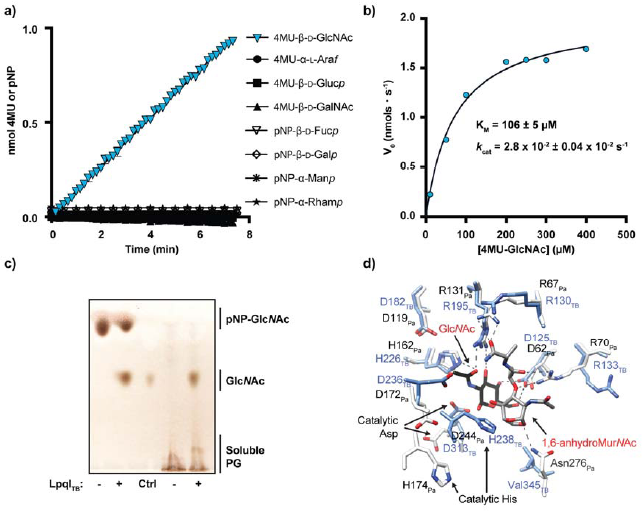
LpqI is an authentic NagZ-type enzyme. **a)** Reactions including 1 μM LpqI and the indicated chromogenic substrates at 1 mM were incubated at 37 °C and release of pNP or 4MU was followed by absorbance or fluorescence respectively. **b)** LpqI was incubated with increasing concentrations of 4MU-Glc*N*Ac. The rate of 4MU release was plotted and the curve fit with the Michaelis-Menton equation using GraphPad Prism 7.0. (n = 3) **c)** LpqI is able to release Glc*N*Ac from soluble muropeptides derived from *M. smegmatis* mc^2^155 PG. **d)** The active site of LpqI is highly conserved as evidenced by the nearly identical positioning of key binding residues observed in the Glc*N*Ac, 1,6-anhydroMur*N*Ac complex with NagZ_Pa_ (PDB: 5G3R) with an overall RMSD of 1.01 Å.

To further confirm its function and validate its role in PG-recycling we solved the 1.96 Å X-ray crystal structure of LpqI (PDB code: 6GFV; Figure S1, Table S3). LpqI consists of a single TIM-barrel domain similar to cytoplasmic Gram-negative orthologs but lacks the C-terminal domain associated with extracellular NagZ enzymes from some Gram-positive bacteria (Figure S3). Alignment of LpqI with the NagZ/Glc*N*Ac/1,6-anhydroMur*N*Ac complex from *Pseudomonas aeruginosa* (NagZ*_Pa_*; PDB:5G3R) or NagZ from *Bacillus subtilis* (PDB:4GYJ) resulted in a root-mean-square deviation of 0.96 Å and 1.01 Å respectively (Figure S3). Superposition of the post-cleavage NagZ*_Pa_* complex with LpqI indicates that the appropriate coordinating residues for Mur*N*Ac or 1,6-anhydroMur*N*Ac recognition are intact in LpqI, supporting its role in PG-recycling (Figure 2d) ^38^.

### LpqI-catalysed utilisation of peptidoglycan components by mycobacteria

Having confirmed the *in vitro* activity of LpqI, we sought to determine the fate of its reaction products, Mur*N*Ac and Glc*N*Ac, in growing *M. bovis* BCG. Prior research has shown that most mycobacteria are unable to use Glc*N*Ac as a sole carbon source, with *M. smegmatis* being one of the notable exceptions^39^. Furthermore, amino acids including L-Ala, L-Glu, and L-Asp have previously been shown to serve as nitrogen sources for *M. tuberculosis* H37Rv^40^. To our knowledge, recycling of Glc*N*Ac or Mur*N*Ac has not been reported, nor has recycling been tested for soluble PG fragments. To evaluate this, *M. bovis* BCG was cultured in minimal media supplemented with glycerol (1% v/v) or Mur*N*Ac (0.2% w/v) in Sauton’s minimal media with constant aeration. As observed in Figure 3a, *M. bovis* BCG was able to grow using Mur*N*Ac as a sole carbon source. To confirm that this was not a phenotype specific to *M. bovis* BCG we also evaluated the ability of *M. tuberculosis* H37Rv to grow on the same carbon sources with identical results (Figure 3b). Intriguingly growth on Mur*N*Ac in broth was heavily dependent on the aeration of the culture. In contrast, growth on glycerol was unaffected by this change (Figure 3c). To further evaluate the potential for mycobacteria to take up Glc*N*Ac but use it for other purposes other than central metabolism, we tested the ability of *M. bovis* BCG to incorporate ^14^C Glc*N*Ac into whole cells. Under different growth conditions (rich medium, carbon-poor medium, aerated cultures, static cultures) we were unable to detect significant amounts of Glc*N*Ac being taken up by *M. bovis* BCG. In all cases the c.p.m. of the label in whole-cells was less than or equal to unlabeled controls. We conclude from these data that pathogenic mycobacteria are able to utilise Mur*N*Ac, but not Glc*N*Ac in an O_2_-dependent fashion.

**Figure 3.**
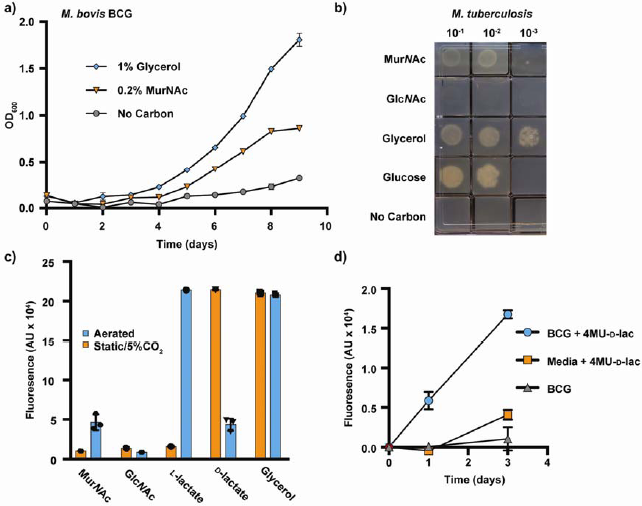
M. tuberculosis and M. bovis BCG are able to recycle Mur NAc. **a)** *M. bovis* BCG WT was inoculated at a starting OD_600_ of 0.1 in minimal media containing glycerol and tyloxopol, Mur*N*Ac or no carbon and growth was monitored daily by taking OD_600_ readings at the indicated concentrations (n = 3). **b)** *M. tuberculosis* H37Rv was washed and then serially diluted into fresh carbon-free minimal media. 10 μL of each dilution was spotted onto Sauton’s agar containing the indicated carbon sources at 5 mM. **c)** Growth of *M. bovis* BCG on 5 mM Mur*N*Ac, Glc*N*Ac, L-lactate, D-lactate, and glycerol was evaluated in aerated or static 5% CO_2_ cultures using a resazurin assay (n = 3). **d)** Mid-exponential *M. bovis* BCG was grown in minimal media with 5 mM glycerol including 1 mM 4MU-D-lactate with constant agitation. At the indicated times the 4MU fluorescence of the samples was determined (n = 3).

### Mechanism of Mur*N*Ac metabolism

Given its structural similarity to Glc*N*Ac, the ability of *M. tuberculosis* and *M. bovis* BCG to grow on Mur*N*Ac was surprising and so we evaluated the biochemical processing steps associated with Mur*N*Ac utilization. Mur*N*Ac is a combination of Glc*N*Ac and D-lactate joined by an ether linkage. This suggests that the bacterium is either using the Glc*N*Ac moiety for glycolysis, or shunting the lactate derived from Mur*N*Ac into the TCA cycle. We tested this by inhibiting glycolysis with 2-deoxyglucose (2DG) in cultures grown using Mur*N*Ac, glucose and glycerol as sole carbon sources (Figure S4). These data suggested that the pathway of Mur*N*Ac utilization did not require glycolysis and indicated that the lactate moiety of Mur*N*Ac, was instead most likely serving as a carbon source. Consistent with this, when used as a sole carbon source, growth on L-lactate and Mur*N*Ac was O_2_ dependent while D-lactate was better utilized under static, 5% CO_2_ culture conditions, where Mur*N*Ac could not be used as a carbon source (Figure 3c).

These data allow us to hypothesize a mechanism by which *M. bovis* BCG metabolises Mur*N*Ac. Given that metabolism of L-lactate and Mur*N*Ac are O_2_-dependent, we anticipate that use of Mur*N*Ac follows cleavage of the D-lactate from Mur*N*Ac *via* an inverting mechanism to produce L-lactate and Glc*N*Ac. In this case, the O_2_-dependency on Mur*N*Ac metabolism is likely the result of an O_2_-dependent lactate monooxygenase. Consistent with this, two O_2_-dependent L-lactate monooxygenases have been identified in *M. tuberculosis* (Rv0694, Rv1872c)^41^. Given the unusual nature of Mur*N*Ac, generation of free lactate by the bacterium would require the activity of a specific lactyl-etherase. To test for the presence of this activity in whole cells, we synthesized a 4MU-D-lactate derivative to serve as a reporter-analog of Mur*N*Ac (Figure S5). Consistent with the presence of a lactyl etherase, cultures of *M. bovis* BCG were able to release 4MU from this compound during the course of growth (Figure 3d). Together these data support a model where mycobacteria cleave the lactyl-moeity from Mur*N*Ac by an as-yet unidentified enzyme and utilise the product of that reaction as a carbon source under aerated conditions.

### LpqI-dependent uptake of PG metabolites by mycobacteria

While our data strongly support metabolism of Mur*N*Ac by *M. bovis* BCG, we wanted to confirm the role of LpqI in mycobacterial PG-recycling. To do this we constructed a mutant strain of *M. bovis* BCG lacking *lpqI* using specialized transduction^16^. To validate that LpqI is the only β-*N*-acetylglucosaminidase produced by *M. bovis* BCG, we used a whole cell β-*N*-acetylglucosaminidase assay. This demonstrated that *M. bovis* BCG Δ*lpqi* is devoid of β-*N*-acetylglucosaminidase activity as the amount of 4MU released is not significantly different from the spontaneous release in sterile media (Figure 4a). This deficiency is complemented by replacement of the *lpqI* gene at a distal chromosomal location under the control of its native promoter (Δ*lpqi*::*lpqI*) and is not complemented by the empty vector (Δ*lpqi*::EV) (Figure 4a). Growth of Δ*lpqi in vitro* is unaltered as compared to the wild-type (Figure 4c). This mutant therefore provided us with an opportunity to probe the role of disaccharide cleavage in mycobacterial PG-recycling.

**Figure 4.**
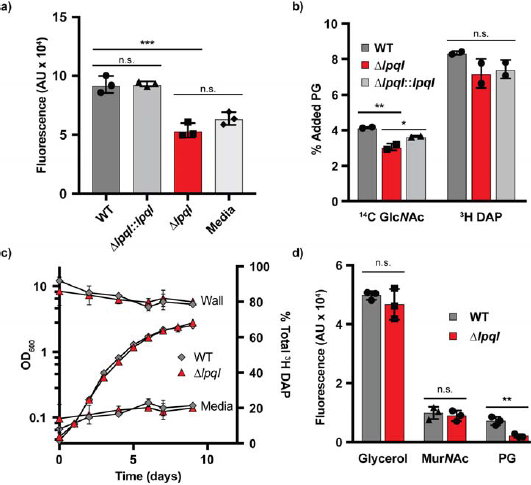
M. bovis BCG is able to recycle PG. **a)** The same strains were incubated with 1 mM 4MU-Glc*N*Ac in minimal media. After 3 days the fluorescence of the cultures were measured (n = 3). **b)** *M. bovis* BCG WT, Δ*lpqI* and Δ*lpqI*::*lpqI* were incubated with 30,000 CPM of ^14^C Glc*N*Ac-labelled muropeptides or 100, 000 CPM of 3H DAP-labelled muropeptides for 10 days after which the cell wall material was isolated and subjected to liquid scintillation counting (n = 2). **c)** *M. bovis* BCG WT and Δ*lpqI* were simultaneously evaluated for release of cell wall peptides and growth (n = 3). **d)** The same strains were evaluated for their growth using glycerol, Mur*N*Ac and PG as sole carbon sources using a resazurin assay (n = 3; *** = p < 0.001; ** = p < 0.005). Statistical significance determined using a two-tailed t-test.

The order in which muropeptides are recycled, and the chemical structure of the recycled material is critical for the immune sensing of the bacterium. To determine the order of PG-recycling steps, we first determined the impact of the loss of *lpqI* on the recycling of cell wall material. To investigate this, we generated radio-labelled muropeptides and tested them in whole-cell uptake assays. Radiolabelled muropeptides had to be generated in *M. smegmatis* due to the inability of *M. bovis* BCG to take-up ^14^C Glc*N*Ac under the conditions we tested. As shown in Figure 4b, the *M. bovis* BCG Δ*lpqI* mutant took up approximately 25% less of the labelled PG than the wild-type (3% vs. 4% respectively). This assay was then repeated using PG fragments labelled with ^3^H-DAP. In this case, the mutant did take up slightly less of the PG than the wild-type, however this difference was not found to be significant using a two-tailed t-test (Figure 4b). To probe this result further, we pre-labelled cells with ^3^H DAP and monitored the release of the label into the culture media. Supporting the data above, we observed no significant differences between the wild-type and the Δ*lpqI* strain with respect to the amount of label released to the media (Figure 4c). From these experiments we concluded that LpqI is involved in amino sugar recovery but is not required for stem-peptide recycling.

Given the inability of *M. bovis* BCG to take up radiolabeled Glc*N*Ac, we were unable to repeat this experiment and follow release of the sugar to the media. To test the impact of deleting *lpqI* on amino sugar recycling by the bacterium, we evaluated its ability to grow on Mur*N*Ac, glycerol and PG. The Δ*lpqI* strain was not deficient for growth on Mur*N*Ac or glycerol, however unlike the wild-type strain it was unable to grow on PG as a sole-carbon source (Figure 4d). Together these data indicate that *in vitro lpqI* is required for PG-derived amino-sugar recycling.

### Phenotypic characterisation of a Δ*lpqI* mutant

As indicated above, loss of LpqI did not alter the growth rate of the bacterium *in vitro.* However, given that NagZ-like proteins have been found to play a role in β-lactam sensitivity in other bacteria we sought to determine the antibiotic sensitivity of the Δ*lpqI* strain. In contrast to inhibition of *P. aeruginosa* NagZ, deletion of *lpqI* resulted in an increase in survival in the presence of lysozyme and all cell-wall active antibiotics tested (Figure 5a-d)^42^. A smaller impact on survival in the presence of the protein synthesis inhibitor chloramphenicol was observed (Figure 5e). This increase in resistance is not due to a change in cell-wall permeability as determined by ethidium bromide uptake (Figure 5f). These data indicate that *lpqI*-dependent amino sugar recycling is important for the expression of antibiotic resistance by mycobacteria *in vitro*.

**Figure 5.**
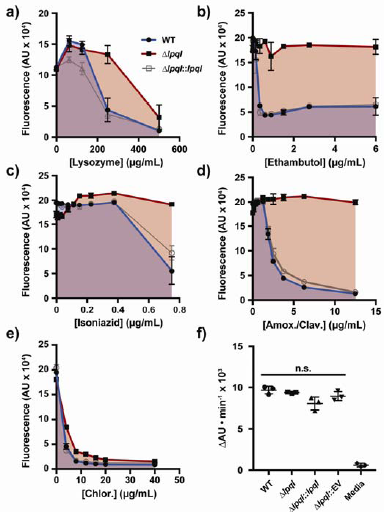
Loss of LpqI leads to lysozyme and antibiotic resistance. **a-e)** *M. bovis* BCG WT, Δ*lpqI* and Δ*lpqI*::*lpqI* were incubated with increasing concentrations of lysozyme or antibiotics at a starting OD_600_ of 0.1. After 7 days incubation total growth was assessed using a resazurin assay, where total fluorescence correlates with respiration and growth (n = 3). f) *M. bovis* BCG WT, Δ*lpqI* and Δ*lpqI*::*lpqI* and Δ*lpqI*::EV were incubated with EtBr and the rate of EtBr was monitored as an increase in fluorescence. No significant differences were found in pairwise t-tests across all strains (n = 3). Statistical significance determined using a two-tailed t-test.

## Discussion

In an attempt to develop diagnostic media for the identification of mycobacteria, several groups in the 1960s observed that *M. tuberculosis* and most other mycobacteria could not metabolise Glc*N*Ac as a sole carbon-source^39,40^. This, along with the absence of known PG recycling-associated genes lead to the assumption that PG recycling is absent in pathogenic mycobacteria. Based on our data and the literature, it is clear that not only is *M. tuberculosis* able to recycle its PG *via* a novel pathway, it is generating two distinct classes of molecules. These are Glc*N*Ac-Mur*N*Ac-peptide, which is sensed by the host, and Glc*N*AcMur*N*Ac which is sensed by the bacterium. While host-sensing of PG is unaffected by the presence of Glc*N*Ac on NOD-stimulatory molecules, our data indicate that LpqI acts as a regulator for Glc*N*Ac-Mur*N*Ac levels by cleaving disaccharides and allowing the break-down of Mur*N*Ac. In other bacteria cell wall damage can trigger various stress responses, and so it is likely that a build-up of Glc*N*Ac-Mur*N*Ac disaccharides may trigger a stress-like response in mycobacteria^43^. Consistent with this, *lpqI* is encoded adjacent to a universal stress response protein in several species of mycobacteria (Figure S5).

As a starting point to investigate PG recycling in *M. tuberculosis*, we investigated the core mycobacterial protein, LpqI. Despite the absence of other known PG-recycling proteins, we have shown that LpqI is an authentic β-*N-*acetylglucosaminidase which is able to cleave PG fragments *in vitro*. Consistent with a role in PG-recycling, *M. bovis* BCG Δ*lpqI* is unable to grow on soluble PG as a sole carbon source, while recycling of the stem-peptide is unaltered in this mutant. Together, our data demonstrate that *M. bovis* BCG and *M. tuberculosis* remove the stem-peptide from PG-fragments prior to disaccharide cleavage and lactyl-ether removal (Figure 6). The processing of Glc*N*Ac-Mur*N*Ac by LpqI prior to lactyl-ether cleavage is also supported by our LpqI crystal structure in which the lactate-binding residue R67 from the *P. aeruginosa* structure is conserved (LpqI: R130) suggesting that the physiological substrate of this enzyme possesses the lactyl group^38^.

**Figure 6.**
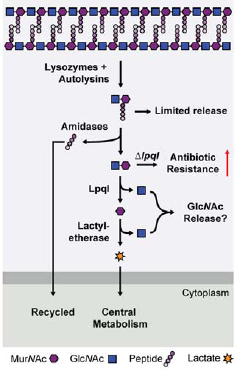
Peptidoglycan recovery pathway in pathogenic mycobacteria. Based on our observations we can propose the following model for PG recycling and recovery in mycobacteria. Cleavage of the cell wall by endogenous autolysins or host-derived lysozyme generates muropeptides. Some of this material undergoes limited release to stimulate the host immune system. The remainder are subsequently degraded by amidases and other peptidases. LpqI then cleaves Glc*N*Ac-Mur*N*Ac, which is followed by D-lactyl-ether cleavage. Lactate can then be used by the cell under aerobic conditions and Glc*N*Ac (or its derivatives) are most likely released. Perturbation of this system by deleting LpqI leads to increased resistance to anti-mycobacterial agents.

Our sole-carbon source assays indicate that while the bacteria are unable to metabolise Glc*N*Ac, surprisingly they can use Mur*N*Ac as a sole carbon source (Figure 3). This is despite the fact that they lack an ortholog of the only known lactyl-etherase, MurQ which cleaves an otherwise stable lactyl-ether in the cytoplasm of most model organisms (Figure 1). Our data indicate that rather than using the Glc*N*Ac portion of the sugar, the bacteria are cleaving the lactyl-ether and metabolising the liberated lactate. During our study we found that *M. bovis* BCG was only able to grow on Mur*N*Ac under aerated conditions. This was also found to be the case for L- but not D-lactate which served as a much better carbon source under O_2_ limiting conditions. As Mur*N*Ac is a combination of D-lactate and Glc*N*Ac, we can predict that the lactyl etherase acting on Mur*N*Ac is proceeding *via* an inverting mechanism. The presence of a specific lactyl-etherase is supported by the turnover of a 4MU-D-lactate reporter compound by *M. bovis* BCG. The O_2_ dependence of this growth is intriguing as Nglycolylation is also an O_2_-dependent activity, suggesting significant alterations to PG metabolism in hypoxic vs. aerobically growing mycobacteria^44^. Consistent with this observation, expression of *lpqI* is upregulated 2-fold during re-aeration after re-activation from non-replicating persistence in the Wayne hypoxia model^45^.

The fate of Glc*N*Ac in this pathway remains unclear, although our data and prior observations suggest that the bacteria do not re-use this sugar. This is surprising given the conservation of the *nagA* (Rv3332) gene in mycobacteria, however it is possible that an alternative pathway exists which involves intermediates not generated under the conditions we have tested. This is hinted at with our ^14^C-labelled muropeptides where incorporation of the labelled-Glc*N*Ac is not expected given the lack of Glc*N*Ac utilisation by the cells. During the production of radio-labelled PG in *M. smegmatis*, a portion of the Glc*N*Ac will have been used by the bacterium to generate Mur*N*Ac rather than strict shunting of Glc*N*Ac into UDPGlc*N*Ac for cell wall biosynthesis. The subsequent removal of the lactyl-ether from this ^14^CMur*N*Ac by *M. bovis* BCG would then follow steps and intermediates we do not yet fully understand. Indeed, bacterial etherases comprise a diverse number of mechanisms and potential reaction products and so a product other than free Glc*N*Ac is entirely possible^46^. We are currently trying to identify and characterise the enzyme responsible for the observed lactyl-etherase activity.

In conclusion, we have identified for the first time a PG recovery pathway in pathogenic mycobacteria. We have shown that this occurs in a step-wise fashion by removing stem-peptide from PG and subsequently cleaving the PG-disaccharide and finally releasing the D-lactate from free Mur*N*Ac. Finally, we have shown that recycling of PG by these bacteria is important for lysozyme and antibiotic resistance.

## Acknowledgements

We wish to thank Sudagar S. Gurcha and Albel Singh for technical assistance. P.J.M. wishes to acknowledge support in the form of a Future Leader Fellowship from the UK Biotechnology and Biological Sciences Research Council (BB/N011945/1). G.V.M. would like to acknowledge funding from UK Biotechnology and Biological Sciences Research Council (BB/P001513/1). A.L.L. acknowledges funding from UK Biotechnology and Biological Sciences Research Council (BB/J015229/1). M.S. would like to acknowledge funding as an Associate FCT-investigator. G.S.B would like to acknowledge support in the form of a Personal Research Chair from Mr James Bardrick and the UK Medical Research Council (grant MR/K012118/1).

## Author Contributions

Conceived of the study: P.J.M. Conceived and designed the experiments: P.J.M., G.V.M., M.S., A.L.L., G.S.B. Performed the experiments: P.J.M., A.R.M., I.T.C, N.V., M.J., M.C. Analysed the data: P.J.M., A.R.M., I.T.C, N.V., M.C. G.V.M., M.S., A.L.L., G.S.B. Wrote and edited the paper: P.J.M, G.V.M., M.S., A.L.L., G.S.B.

